# chromstaR: Tracking combinatorial chromatin state dynamics in space and time

**DOI:** 10.1101/038612

**Authors:** Aaron Taudt, Minh Anh Nguyen, Matthias Heinig, Frank Johannes, Maria Colomé-Tatché

## Abstract

**Background:** Post-translational modifications of histone residue tails are an important component of genome regulation. It is becoming increasingly clear that the combinatorial presence and absence of various modifications define discrete chromatin states which determine the functional properties of a locus. An emerging experimental goal is to track changes in chromatin state maps across different conditions, such as experimental treatments, cell-types or developmental time points.

**Results:** Here we present chromstaR, an algorithm for the computational inference of combinatorial chromatin state dynamics across an arbitrary number of conditions. ChromstaR uses a multivariate Hidden Markov Model to determine the number of discrete combinatorial chromatin states using multiple ChIP-seq experiments as input and assigns every genomic region to a state based on the presence/absence of each modification in every condition. We demonstrate the advantages of chromstaR in the context of three common experimental data scenarios. First, we study how different histone modifications combine to form combinatorial chromatin states in a single tissue. Second, we infer genome-wide patterns of combinatorial state differences between two cell types or conditions. Finally, we study the dynamics of combinatorial chromatin states during tissue differentiation involving up to six differentiation points. Our findings reveal a striking sparcity in the combinatorial organization and temporal dynamics of chromatin state maps.

**Conclusions:** chromstaR is a versatile computational tool that facilitates a deeper biological understanding of chromatin organization and dynamics. The algorithm is implemented as an R-package and freely available from http://bioconductor.org/packages/chromstaR/.

## Introduction

Epigenetic marks such as DNA methylation or histone modifications play a central role in genome regulation. They are involved in a diversity of biological processes such as lineage commitment during development [1], maintenance of cellular identity [2, 3] and silencing of transposable elements [4]. The modification status of many histone marks has been extensively studied in recent years, first with ChlP-chip and later with ChIP-seq, now the de-facto standard procedure for genome wide mapping of protein-DNA interactions and histone modifications. Since its advent in 2007 [1, 2, 5], ChIP-seq technologies have been widely used to survey genome-wide patterns of histone modifications in a variety of organisms [6, 7, 2], cell lines [8] and tissues [9, 10].

The multitude of possible histone modifications has led to the idea of a “histone code” [11, 12], a layer of epigenetic information that is encoded by combinatorial patterns of histone modification states (Fig. 1a). Major resources have been allocated in recent years to decipher this code, culminating in projects such as the ENCODE [13] and Epigenomics Roadmap [10]. Following their examples, most experiments nowadays are designed to probe several histone modifications at once, and often in various cell types, strains and at different developmental time points. These types of experiments pose new computational challenges, since initial solutions were designed to analyze one modification and condition at a time, therefore treating them as independent. Indeed, a commonly used strategy has been to perform peak calling for each experiment separately (univariate analysis) and to combine the peak calls post-hoc into combinatorial patterns [14, 15]. This approach is problematic for several reasons: Because of the noise associated with ChIP-seq experiments and peak calling, combining univariate peak calls will lead to the discovery of spurious combinatorial states that do not actually occur in the genome. Furthermore, different tools or parameter settings are often used for different modifications (e.g. peak calling for broad or narrow marks), making the outcome sensitive to parameter changes and control of the overall false discovery rate difficult. Lastly, this approach requires ample time and bioinformatic expertise, rendering it impractical for many experimentalists.

**Figure 1.**
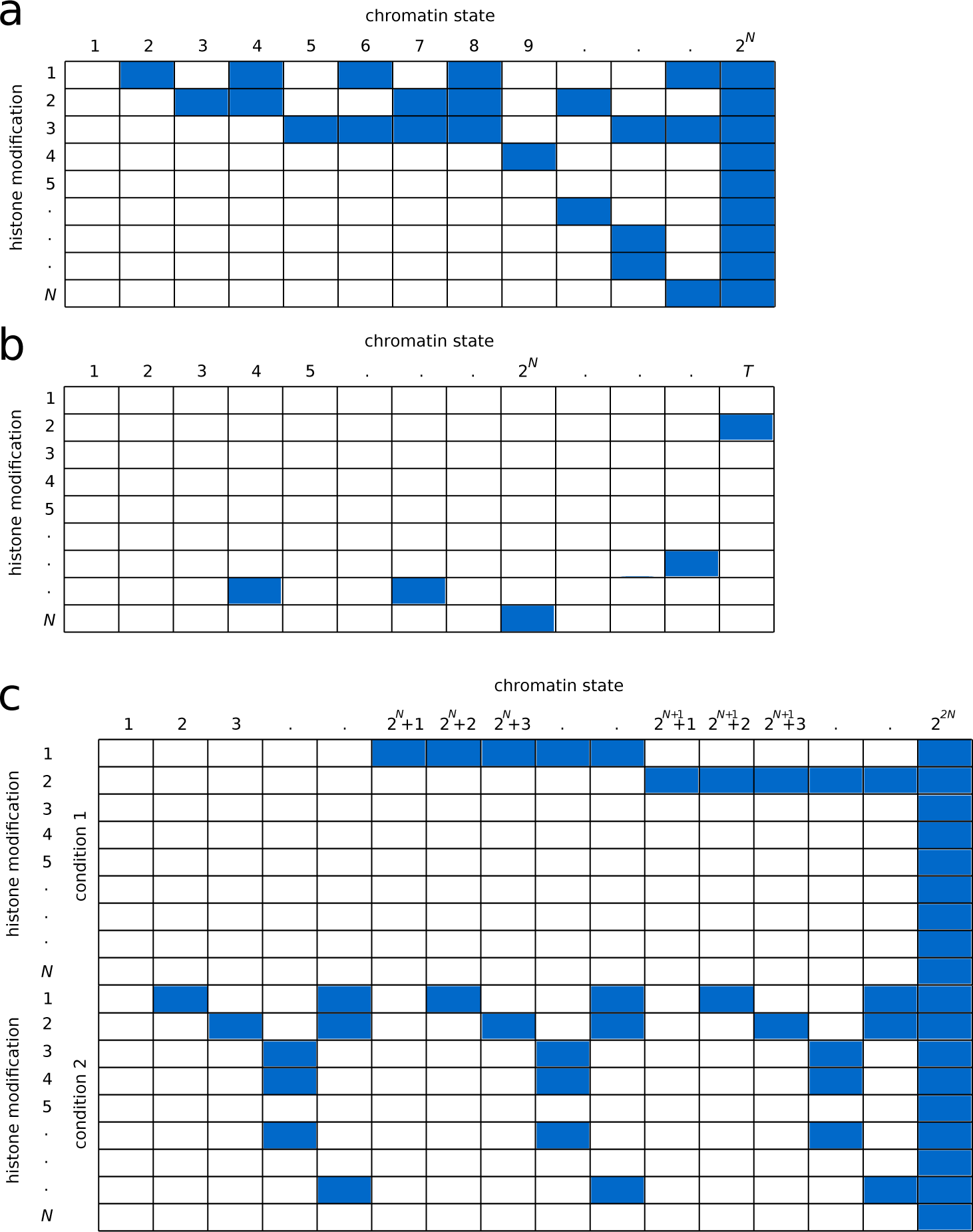
Definition of chromatin states. **(a)** Combinatorial chromatin state definition: Based on the presence (blue) or absence (white) of a histone modification, a chromatin state is the combination of the presence/absence calls at a given position. With *N* histone modifications there are 2^*N*^ different chromatin states. **(b)** Probabilistic chromatin state definition: Each chromatin state has a probability (shades of blue) of finding a histone modification at a given position. Note that a probabilistic state can consist of multiple combinatorial states and vice versa. There is in principle no upper limit for the number of possible probabilistic chromatin states (here, *T*). **(c)** Differential combinatorial chromatin states across two conditions: Based on the presence (blue) or absence (white) of a histone modification across different conditions. With *N* histone modifications and *M* conditions there are 2*^N×M^* different states.

Accurate inferences regarding combinatorial histone modification patterns are necessary to be able to understand the basic principles of chromatin organization and its role in determining gene expression programs. One way forward is to develop computational algorithms that can analyze all measured histone modifications at once (i.e. combinatorial analysis) and across different conditions (i.e. differential analysis). Some methods have been designed to integrate histone modifications into unified chromatin maps [16, 17, 18, 19, 20, 21, 22]. These methods can be classified into three different categories [23]: combinatorial (which define chromatin states based on the presence/absence of every histone modification) [19], continuous (which define chromatin states based on the shape of the ChIP-seq signal) [24, 25], and probabilistic (which have probabilities associated with finding each mark in a given state) [16, 17, 18, 20, 21, 22]. A major drawback of the majority of these approaches [16, 17, 18, 20, 21, 22] is the need to specify the number of distinct chromatin states beforehand, which is usually not known *a priori*. Moreover, in the probabilistic interpretation the inferred states can consist of multiple and overlapping combinatorial states (Fig. 1b). This probabilistic state definition is useful to reduce noise and to identify functionally similar genomic regions for the purpose of annotation, but at the same time it obscures a more direct interpretation of combinatorial states in terms of the presence/absence patterns of the underlying histone modifications.

Finally, none of these methods is designed for comparing chromatin maps across conditions. ChromDiff [26] and dPCA [27] are comparative methods that identify significant chromatin differences between groups of samples. ChromDiff discovers groups of epigenomic features which are the most discriminative in group-wise comparisons of samples, while dPCA uses a small number of differential principle components to explore differential chromatin patterns between two groups of samples. Both methods are useful for identifying defining features of each group, however they do not provide complete information about the genome-wide localization of all chromatin differences between all samples.

In order to overcome these problems we have developed chromstaR, a method for multivariate peak- and broad-region calling. chromstaR has the following conceptual advantages: 1) Every genomic region is assigned to a discrete, readily interpretable combinatorial chromatin state, based on presence/absence of every histone mark, providing a mechanistic interpretation of chromatin states which allows for better insights into how they regulate genome function. 2) The number of chromatin states does not have to be preselected but is a result of the analysis. 3) Histone modifications with narrow and broad profiles can be combined in a joint analysis along with an arbitrary number of conditions. 4) The same approach can be used for mapping combinatorial chromatin states in one condition, or for identifying differentially enriched regions between several conditions, or for both situations combined. 5) Our formalism offers an elegant way to include replicates as separate experiments without prior merging.

The algorithm is implemented in C++ and available as R package to combine speed and ease-of-use, and offers functions to perform the most frequent downstream analysis tasks.

We demonstrate the advantages of chromstaR in the context of three common experimental scenarios (Fig. S1b). First, we consider that several histone modifications have been collected on a single tissue at a given time point (Fig. S1b, Application 1). The goal is to infer how these different modifications combine to form distinct combinatorial chromatin states and to describe their genome-wide distribution. Second, we consider that several histone modifications have been collected in two different cell types or conditions (Fig. S1b, Application 2). Here, the goal is to infer genome-wide patterns of combinatorial state differences between cell types or conditions. Third, we consider the more complex secenario where several histone modifications have been collected for multiple different time points or tissue types (Fig. S1b, Application 3). In this case, the goal is to infer how combinatorial chromatin states are modified during tissue differentiation or development. These three experimental scenarios broadly summarize many of the data problems that biologists and bioinformaticians currently face when analyzing epigenomic data. We show that chromstaR provides biologically meaningful results to these types of data problems, and facilitates deeper biological insights into the dynamic coordination of combinatorial chromatin states in genome regulation.

## Results

### Brief overview of analytical approach and validation of peak calls

Consider *N* ChIP-seq experiments: *N* histone modifications measured in one condition, or one histone modification measured in *N* conditions, or a combination of the two. After mapping the sequencing reads to the reference genome our method consists of two parts (Fig. 2), a univariate peak calling step to estimate the distribution parameters, and a multivariate peak calling step to integrate information from all experiments:

**Figure 2.**
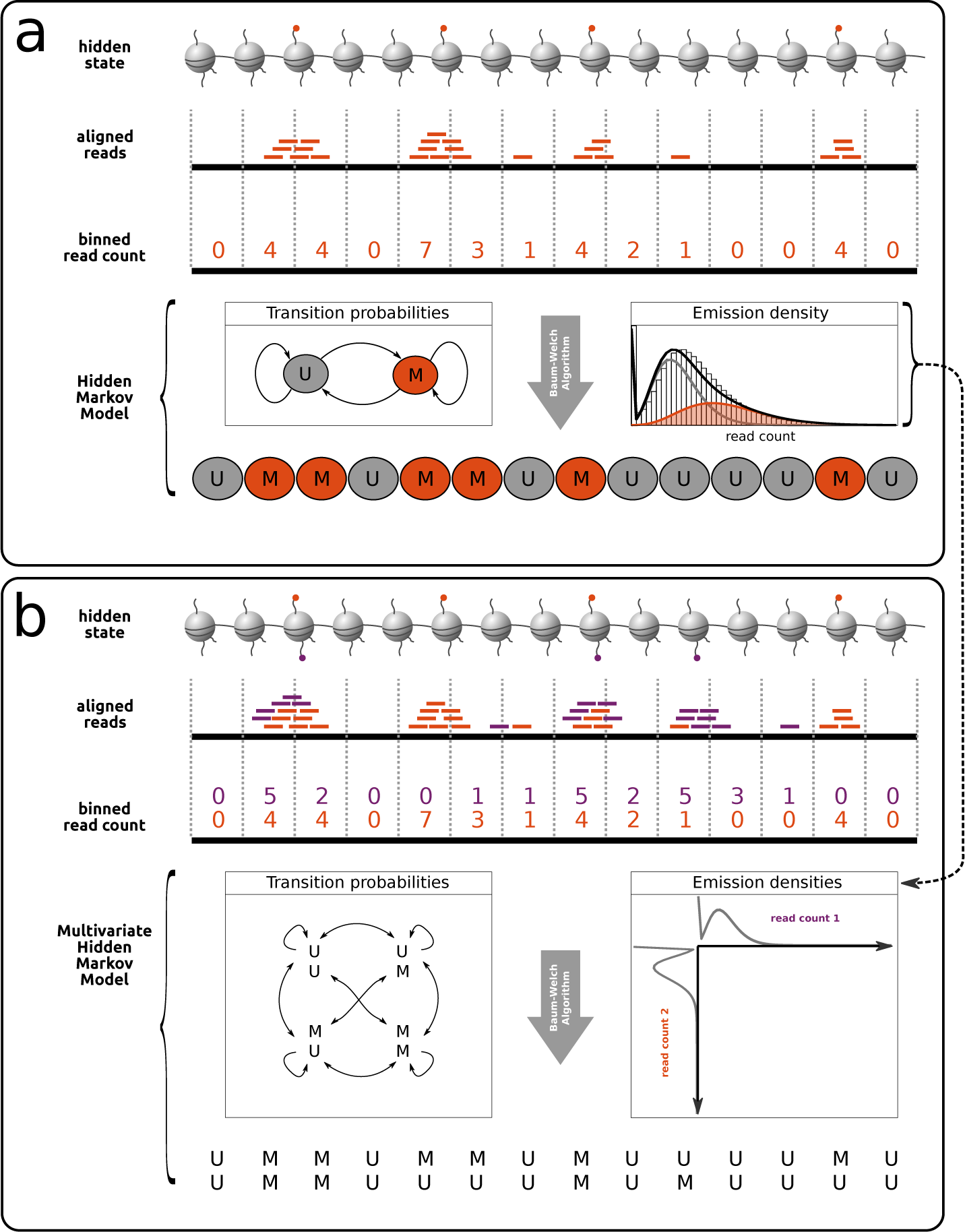
Overview of analytical approach. **(a)** Univariate peak calling: Aligned reads are counted in equidistant, non-overlapping bins. The resulting read count serves as observable for a Hidden Markov Model with components “unmodified” (U) and “modified” (M). Parameters are fitted with the Baum-Welch algorithm. **(b)** Multivariate peak calling: A multivariate emission density is constructed from the densities of step (a), illustrated here for two dimensions. With *N* ChIP-seq experiments, the resulting multivariate density has 2*^N^* components. For the two illustrated dimensions (ChIP-seq 1 and 2), univariate densities are plotted on the x- and y-axis, and the 4 components of the multivariate density are indicated by shaded areas, corresponding to both unmodified (UU - gray), both modified (MM - blue) and unmodified/modified (MU - yellow, UM - green).

(1) Univariate peak calling (Fig. 2a): For each ChIP-seq experiment, we partition the genome into non-overlapping bins (default 1kb) and count the number of reads that map into each bin (i.e. the read count) [28]. We model the read count distribution as a two-component mixture of zero-inflated negative binomials [29, 30], with one component at low number of reads that describes the background noise and one component at high number of reads describing the signal. We use a univariate Hidden Markov Model (HMM) with two hidden states (i.e. unmodified, modified) to fit the parameters of these distributions [31].

(2) Multivariate peak calling (Fig. 2b): We consider all ChIP-seq experiments at once and assume that the multivariate vector of read counts is described by a multivariate distribution which is a mixture of *2^N^* components. We use a multivariate HMM to assign every bin in the genome to one of the multivariate components. The multivariate emission densities of the multivariate HMM, with marginals equal to the univariate distributions from step (1), are defined using a Gaussian copula [32]. A detailed description can be found in **Methods**.

The univariate part of our model (step 1 above), which serves as the basis for the construction of the multivariate model, provides high-quality peak calls that measure up against existing methods. We compared our method with two commonly used peak callers, Macs2 [33] and Sicer [34], using publicly available datasets of qPCR validated regions [35]. We compared the performance of the three methods on two datasets, one for H3K4me3 (narrow profile), and one for H3K27me3 (broad profile). The H3K4me3 dataset had 33 qPCR validated regions and the H3K27me3 dataset had 197 qPCR validated regions. The ChIP-seq datasets were analyzed with the standard settings of each peak caller (**SI text**), and each base pair was assigned a score by the algorithm. This output was used to compute receiver operator characteristic (ROC) curves and area-under-curve (AUC) values [36]. The performance of chromstaR for these datasets in terms of the AUC is equal or better than that of Macs2 and Sicer (Fig. S2). For the multivariate peak calling, in the absence of datasets with validated peak calls for multiple marks, we performed a simulation study based on parameters obtained from real data (**SI text**). ROC curves and AUC values show that multivariate peak calls are of high quality (Fig. S3).

### Application 1: Mapping combinatorial chromatin states in a reference tissue

Lara-Astiaso et al. [37] measured four histone modifications (H3K4me1, H3K4me2, H3K4me3 and H3K27ac) and gene expression in 16 mouse hematopoietic cell lines and their progenitors (Fig. S1). All four marks have a relatively narrow ChIP-seq profile. The authors’ goal was to document the dynamic enhancer landscape during hematopoietic differentiation. With four measured histone modifications there are 2^4^ = 16 possible combinatorial states defined by the presence/absence of each of the modifications. In order to provide a snapshot of the genome-wide distribution of these combinatorial states in a given cell-type, we applied chromstaR to the ChIP-seq samples collected from monocytes (see Fig. S6 for the analysis of other cell types). In the following we introduce a shorthand notation where combinatorial states are denoted between brackets [ ] and each mark is abbreviated by its chemical modification. For example, the combination [H3K4me1+H3K4me2+H3K27ac] will be abbreviated as [me1/2+ac]. If we use the full name of a mark (e.g. “H3K4me1”) we are referring to the mark in a classical, non-combinatorial, context. See Fig. 3d for all combinations with shorthands.

**Figure 3.**
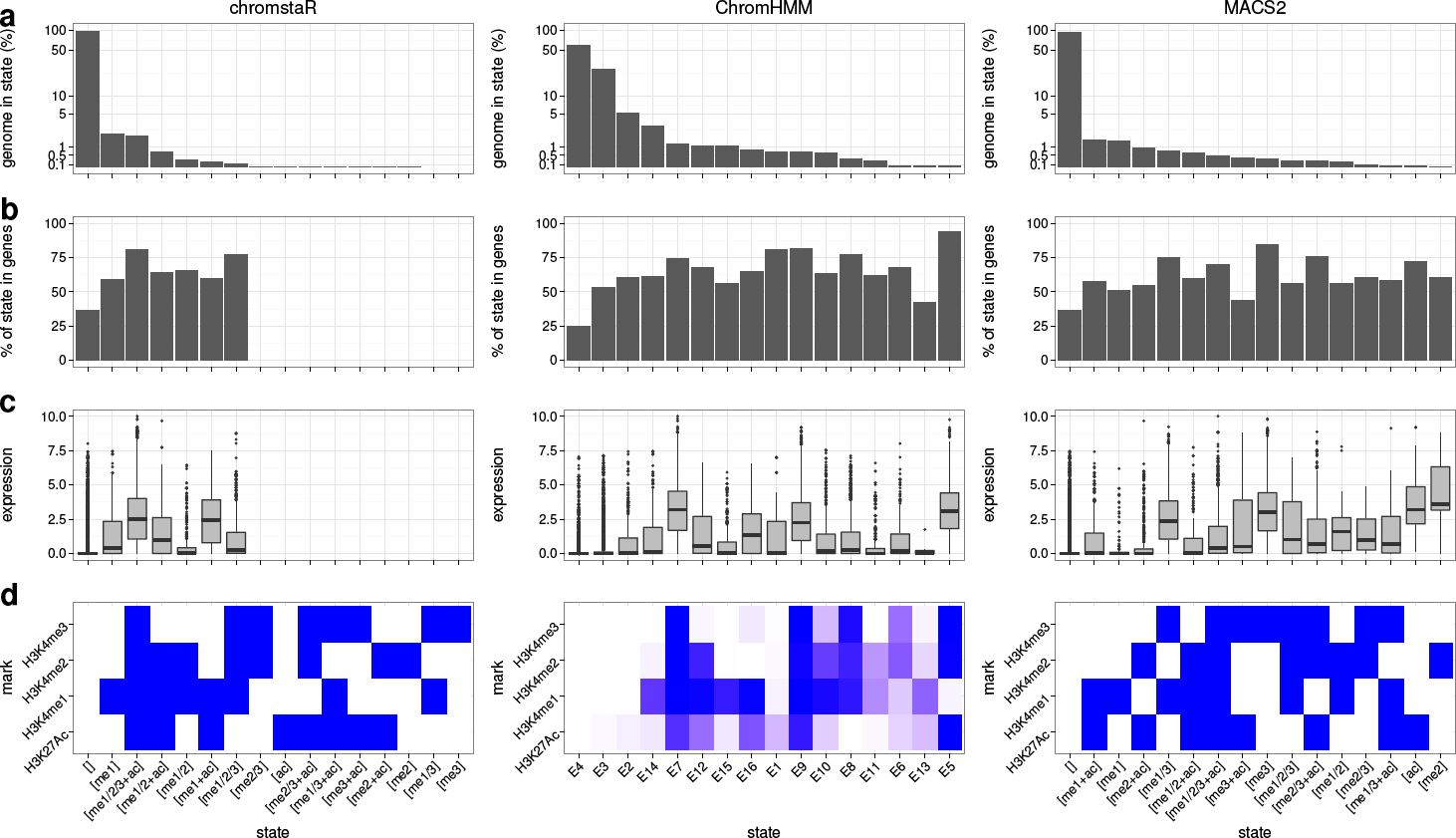
Chromatin states in monocytes. **(a)** Genomic frequency, i.e. the percentage of the genome that is covered by the chromatin state. The sum over all states equals 100%. **(b)** Overlap with known genes. **(c)** Expression levels of genes whose transcription start site (TSS) overlaps the chromatin state. **(d)** Heatmap showing the chromatin state definition. Histones in chromstaR and MACS2 states are either present (blue) or absent (white). ChromHMM states have a continuous emission probability from zero (white) to one (blue).

chromstaR found that many of the 16 possible combinatorial states were nearly absent at the genome-wide scale, with 7 of the 16 states accounting for nearly 100% (>99.99%) of the genome (Fig. 3a). This observation indicates that the “histone code” defined by these four histone modifications is much less complex than theoretically possible, perhaps as a result of biochemical constraints on the co-occurance of certain modifications on the same or neighboring aminoacid residues. However, some of the discovered chromatin states display “incompatible” combinations (the ones displaying more than two modifications on the same histone and residue, such as for example [me1/2/3]). Re-analysis of the data finds eight of the 16 states present in the genome, with a smaller frequency of incompatible states (Fig. S4). These results show that these states are in part due to having pooled data from several nucleosomes into the same bin, but are probably also caused by antibody cross-reactivity and residual cell heterogeneity.

The empty state [ ], which we here define as the simultaneous absence of all measured marks at a given genomic position, was the most frequent state, covering 94.8% of the genome. The high prevalence of this state reflects the fact that Lara-Astiaso et al. [37] focused on marks with a narrow profile that had previously been shown to occur proximal to genic sequences [38, 2, 3]. Indeed, only 36% of the empty state overlapped known genes while the remaining 64% mapped to nongenic regions throuhgout the genome, and probably tag other (unmeasured) histone modifications, such as repressive heterochromatin-associated marks.

In order to evaluate chromatin state frequencies on a data set with a mixture of broad and narrow histone modifications, we analyzed human Hippocampus tissue data from the Epigenomics Roadmap with seven marks [10] and IMR90 cell line data from the ENCODE project with 26 marks [13] (see **Methods**). In the Hippocampus data we found that only 21 out of the 128 possible combinatorial states were necessary to explain more than 99% of the epigenome, and indeed the empty state covered only 32% of the genome (Fig. S7). Moreover, in the lung fibroblast cell line we found that from the possible 67 million states only 0.02% (~12000) are needed to explain more than 95% of the genome, while the empty state covered only 16% of it, showing that when more marks are included the percentage of the genome in the empty state decreases [23].

Contrary to the empty state, on average 68% (range: 59-81%) of the genomic regions found to be in one of the 6 most frequent (non-empty) combinatorial states in mouse monocytes overlap known genes (Fig. 3b), thus suggesting an active role in the regulation of gene expression. To assess this, we examined the combinatorial state profiles of the 6 most frequent states relative to the transcription start site (TSS) of expressed and non-expressed genes (Fig. 4a). In contrast to non-expressed genes, expressed genes were clearly characterized by the presence of state [me1/2/3+ac] proximal to the TSS. This is consistent with previous reports that have used H3K4me3 together with H3K27ac to tag active promoters [39]. However, our analysis also uncovered a more subtle enrichment of state [me1] shouldering the TSS (Fig. 4a). We found that 18% of [me1] sites occur in regions directly flanking state [me1/2/3+ac] and 61% of all [me1] can be found within 10kb of [me1/2/3+ac] sites (see Fig. 5 for an example). These two states therefore constitute a single, broad chromatin signature that defines a subset of expressed genes. Interestingly, this subset of genes had significantly higher expression levels (*p* ≈ 10^−101^, Wilcoxon rank-sum test) and distinct GO terms compared with genes marked only by the active promoter state (i.e. [me1/2/3+ac] at the TSS and no [me1] in flanking regions, Fig. 6 and Table 1). This observation suggests that the co-occurance of [me1/2/3+ac] and [me1] in broad regions surrounding the TSS marks what may be called “enhanced” active promoters ([me1/2/3+ac] + [me1]).

**Figure 4.**
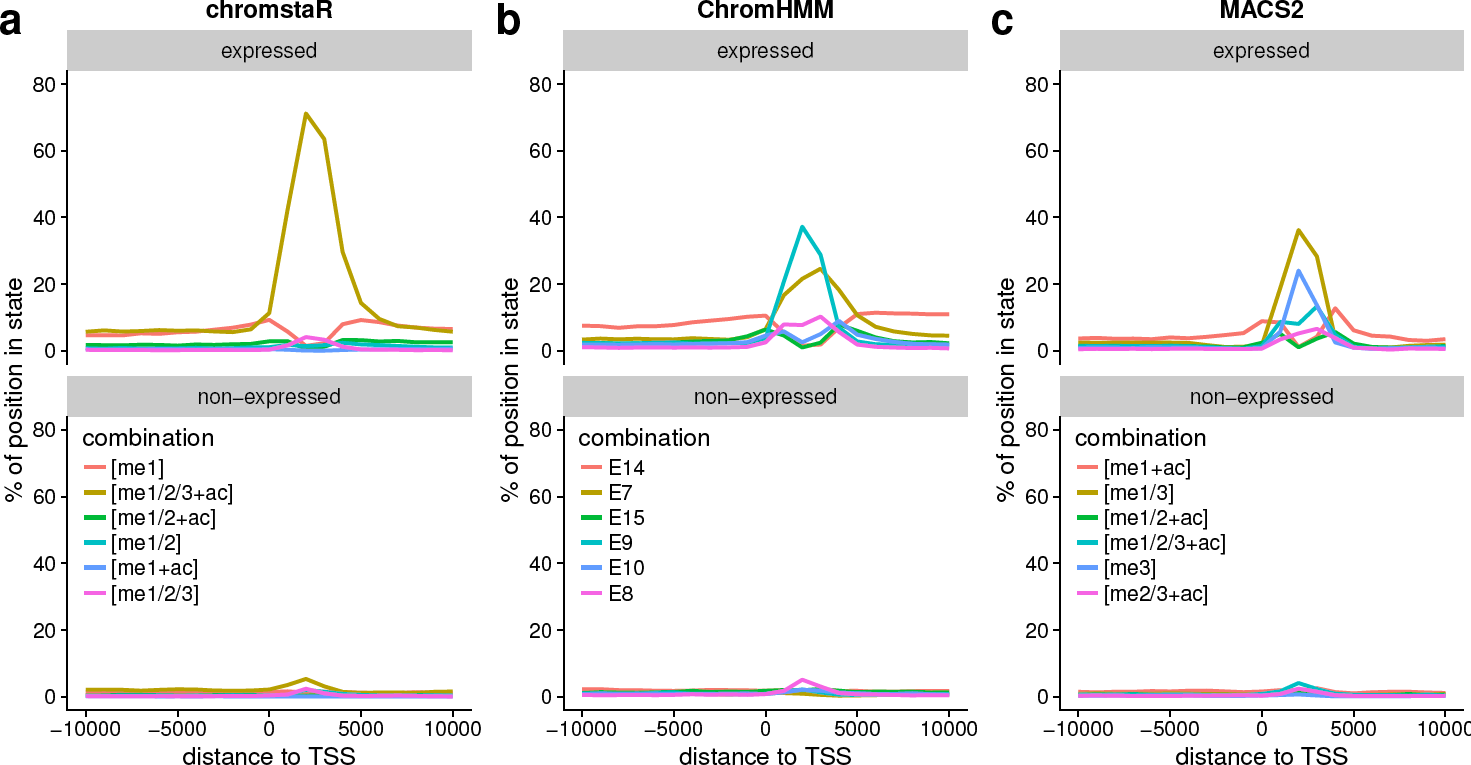
Fold enrichment of chromatin states around transcription start sites (TSS) of expressed (top) and non-expressed (bottom) genes. Shown are the enrichment profiles for the 6 states that are most enriched around TSS. chromstaR consistently assigns state [me1/2/3+ac] to expressed TSS and has a higher sensitivity for detecting those, while the other two methods assign two states with lower sensitivity.

**Table 1.** The first 10 significant gene ontology terms for TSS overlapping the [me1/2/3+ac] state with the [me1] state flanking it, versus the TSS overlapping the [me1/2/3+ac] state. Numbers indicate the binomial false discovery rate (BinomFdrQ) as reported by GREAT.

|  | [me1/2/3+ac] + flanking [me1] |  | [me1/2/3+ac] |  |
| --- | --- | --- | --- | --- |
| 1 | posttranscriptional regulation of gene expression | 5.75e-25 | RNA processing | 1.40e-36 |
| 2 | regulation of translation | 1.57e-21 | ncRNA metabolic process | 6.75e-34 |
| 3 | peptidyl-lysine modification | 1.69e-17 | ncRNA processing | 2.57e-29 |
| 4 | microtubule nucleation | 2.80e-13 | DNA repair | 1.14e-25 |
| 5 | mRNA transport | 1.23e-10 | ribosome biogenesis | 3.36e-17 |
| 6 | RNA localization | 3.61e-10 | rRNA metabolic process | 9.14e-17 |
| 7 | RNA transport | 7.01e-10 | tRNA metabolic process | 3.14e-16 |
| 8 | GPI anchor biosynthetic process | 1.01e-09 | rRNA processing | 1.85e-15 |
| 9 | negative regulation of translation | 1.16e-09 | protein folding | 3.74e-14 |
| 10 | DNA replication | 1.60e-09 | tRNA processing | 1.02e-11 |

To compare the results obtained with chromstaR to other computational approaches, we first re-analyzed replicate datasets from the mouse monocytes [37], the human Hippocampus [10] and the lung fibroblast cell line data [13] using several publicly available methods [16, 19, 21, 22] (**SI text**). chromstaR is the method that provides the best performance in assigning a consistent segmentation between replicates (Fig. S5c) and in detecting regions with high read count fold change as differential (Fig. S5d). Among the alternative methods, ChromHMM provides the best performance and flexibility [16], we will therefore use it in the following for comparison purposes. ChromHMM employs a multivariate HMM to classify the genome into a preselected number of probabilistic chromatin states, and was used to annotate the epigenome in the ENCODE [13] and Epigenomics Roadmap [10] projects. It therefore offers a method to segment the genome into a set of probabilistic chromatin states that can then *a posteriori* be interpreted at the biological level. We also compare the results obtained with chromstaR to the ones obtained using MACS2 [33], because it is one of the most widely used univariate peak callers.

**Figure 5.**
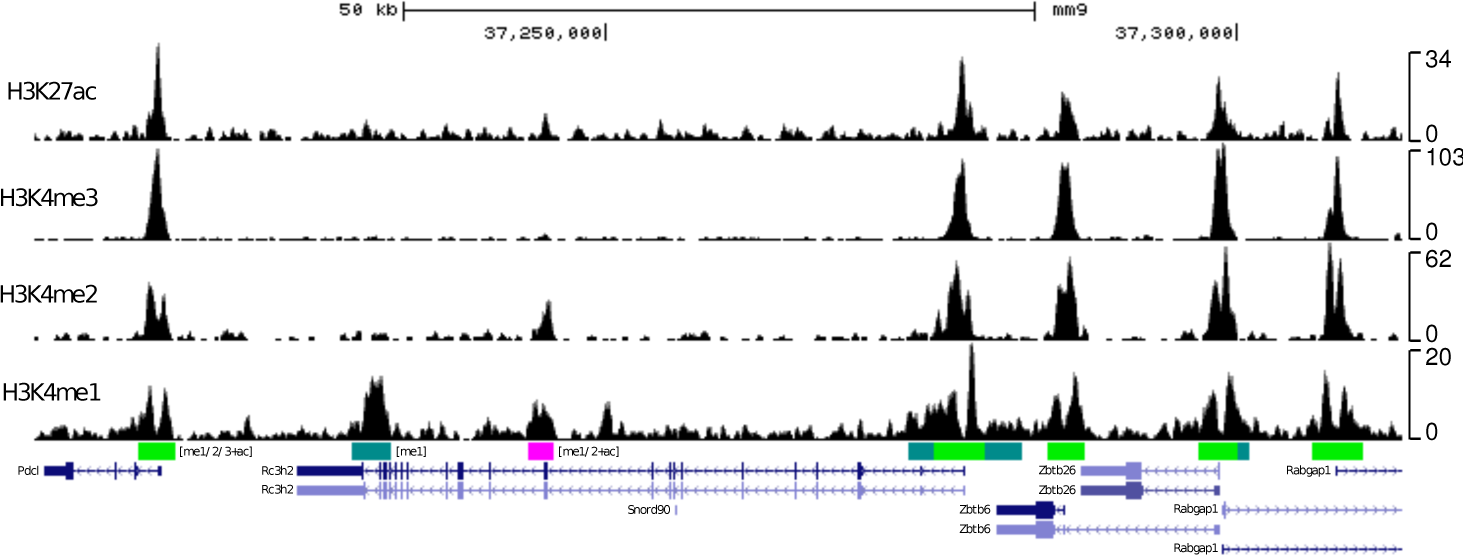
Genome browser snapshot showing a 100kb region on chromosome 2 with several active promoter signatures [me1/2/3+ac] (green) flanked by the [me1] signature (teal). The four black tracks show normalized signal coverage profiles for the four measured histone modifications. The colored track below shows combinatorial state calls, and the bottom track shows genes. Promoter signatures coincide with gene starts.

When using a multivariate segmentation method like ChromHMM, the number of chromatin states needs to be decided beforehand, which is difficult as this number is rarely known *a priori*. In the absence of detailed guidelines we fitted a 16 state model to the mouse hematopoietic data. Our comparison uncovered substantial method-specific differences in state frequencies (Fig. 3). Both ChromHMM and MACS2 found all 16 states present in the genome with more than 0.01% genome coverage. To understand how state-calls compared between methods, we evaluated to which extent the states detected by one method coincided with those detected by the other method(s) (Fig. S8). Most notable, we found that genomic regions corresponding to chromstaR’s active promoter state [me1/2/3+ac] were assigned to two alternative states (E7 and E9) by ChromHMM. These latter two states were very similar in terms of their emission densities, but significantly different at the level of gene expression (*p* ≈ 10^−90^, t-test, Fig. 3c). Moreover, chromstaR’s single empty state [ ] corresponded to two functionally similar (nearly) empty states (E3, E4) detected by ChromHMM. A third almost empty state E2 with weak H3K27ac signal had slightly higher expression levels than the other two empty states and also overlapped with chromstaR’s empty state [ ] (Fig. S8b). These state redundancies highlight the difficulty in selecting the number of chromatin states for ChromHMM, for without extensive manual curation it is difficult to know if two states are truly redundant (likely E3 and E4) or if they are biologically different on some level (E7 and E9).

Although MACS2 is not designed for multivariate analysis, we constructed *ad hoc* combinatorial state calls from the univariate analyses obtained from each ChIP-seq experiment to illustrate the problems of this commonly used analysis technique. As expected, MACS2 results were noisy: many of the combinatorial states detected by chromstaR showed very heterogenous state calls with MACS2 (Fig. S8a). For instance, a considerable proportion (35%) of genomic regions detected by chromstaR as being in the active promoter state [me1/2/3+ac] were assigned to another promoter state (containing H3K4me3) by MACS2. We suspect that this is due to the limitations of MACS2 in calling broader marks (e.g. H3K4me1) or moderate enrichment with the default parameters, which results in frequent missed calls for individual modifications, and subsequently also in the limited detection of ‘complex’ combinatorial states such as [me1/2/3+ac] that are defined by the presence of all modifications.

To better understand the functional implications of the state frequency and state pattern differences between these methods, we evalute the chromatin state signatures of both ChromHMM and MACS2 around TSS of expressed and non-expressed genes (Fig. 4b,c). In contrast to chromstaR, chromatin signatures obtained by the other two methods did not as effectively distinguish these two classes of genes, suggesting that chromstaR has a higher sensitivity for detecting these signatures (Table 2).

**Table 2.** Performance for detecting expressed TSS.

|  | Sensitivity | Precision | F1-score |
| --- | --- | --- | --- |
| chromstaR | 0.71 | 0.97 | 0.82 |
| Macs2 | 0.60 | 0.98 | 0.75 |
| ChromHMM | 0.59 | 0.98 | 0.73 |

### Application 2: Differential analysis of combinatorial chromatin states

In order to understand combinatorial chromatin state signatures that are specific to a given cell type or disease state, it is necessary to compare at least two different tissues with each other, or a case and a control. In this context, the goal is to identify genomic regions showing differential (or non-differential) combinatorial state patterns. Such differential patterns are indicative of regions that underly the tissue differences and are therefore of substantial biological or clinical interest. chromstaR solves this problem by considering all *2^2N^* possible combinatorial/differential chromatin states (Fig. 1c), where *N* is the number of histone modifications measured in both conditions. Out of the 2*^2N^* states, 2*^N^* are non-differential and 2*^2N^* − 2*^N^* are differential.

We anlayzed two differentiated mouse hematopoietic cells (monocytes versus CD4 T-cells) from [37], with four histone marks each (H3K4me1, H3K4me2, H3K4me3 and H3K27ac). We found that 5.37% of the genome showed differences in combinatorial state patterns between the two cell types (Fig. 7a, example browser shot in Fig. 8). The most frequent differential regions involved the [me1] combination (2.37%) followed by regions with the [me1/2/3+ac] combination (0.92%). These differences are even more striking when viewed in relative numbers: 59% of the [me1/2/3+ac] sites were concordant between the two cell types, while only 8% of the [me1] sites were concordant. This is in line with previous findings showing that H3K4me1 is highly cell type specific [40, 41, 42, 43].

In order to determine if these differences in chromatin play a role in cellular identity, we explored gene expression differences for differential chromatin states. We found that loss of state [me1] as well as of state [me1/2/3+ac] is correlated with a decrease in expression levels (Fig. 7b). This is consistent with our previous observation (section **Application 1**) that [me1/2/3+ac] defines active promoters and [me1] together with [me1/2/3+ac] defines enhanced active promoters (Fig. 6). To investigate the function of the differential loci, we performed a GO term enrichment of these regions [44] and found an impressive confirmation of cell type identity in the GO terms (Table S1): While regions that are marked by [me1/2/3+ac] or [me1] in both cell types show enrichment for general immune cell differentiation terms, regions that are marked with [me1] or [me1/2/3+ac] only in CD4 T-cells show terms such as “T-cell activation and differentiation”. Vice versa, regions that are marked with those signatures in monocytes but not in T-cells show enrichment of terms such as “response to other organism” and “inflammatory response”.

**Figure 6.**
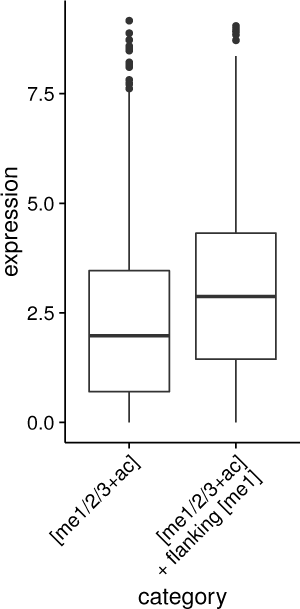
Expression levels of genes whose transcription start site (TSS) shows either the [me1/2/3+ac] signature alone or the [me1/2/3+ac] signature flanked by [me1]. TSS flanked by [me1] show significantly higher expression levels (*p* ≈ 10^−101^, Wilcoxon rank-sum test).

**Figure 7.**
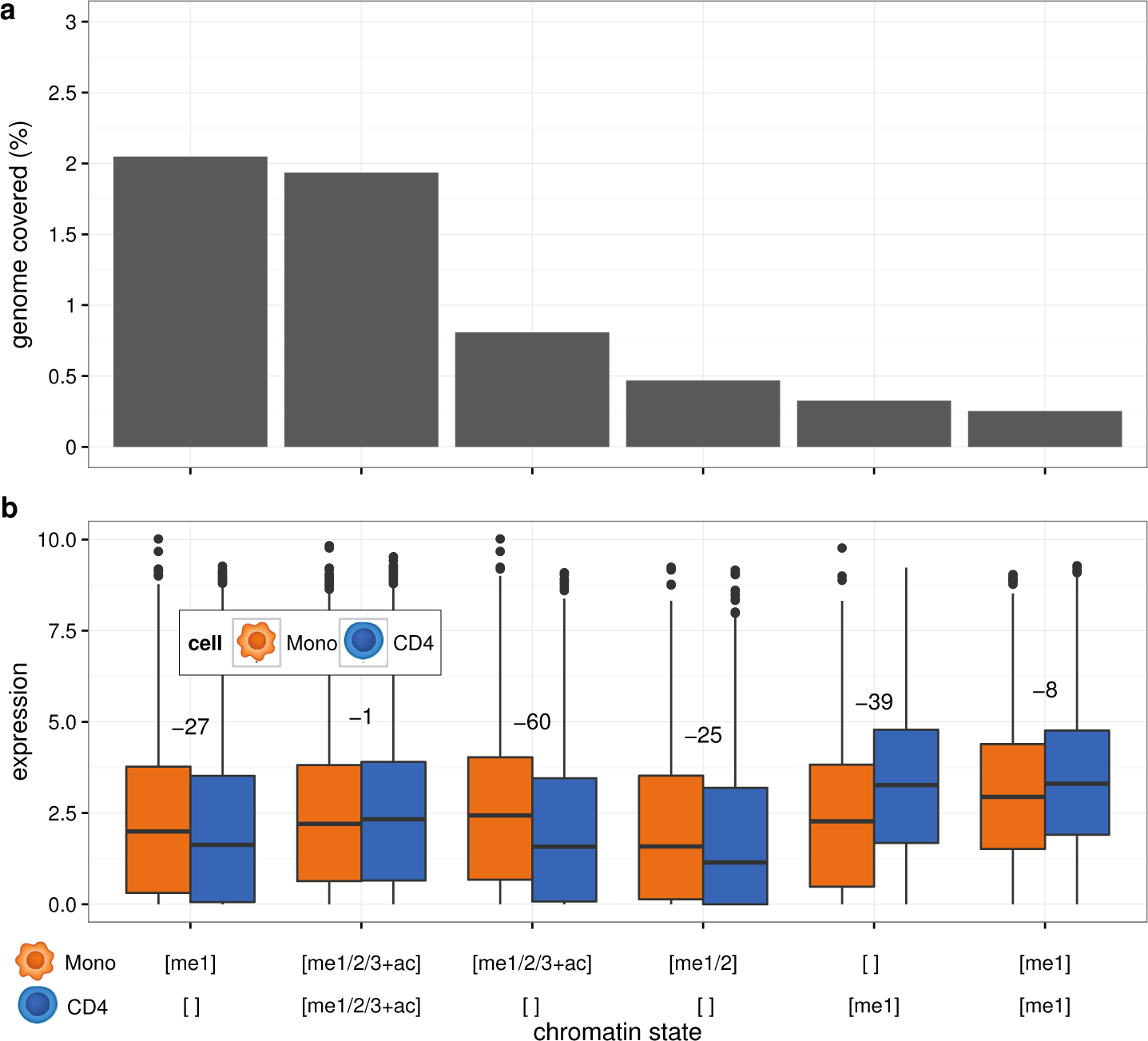
Differential analysis of monocytes and CD4 T-cells. **(a)** Genomic frequency of the six most frequent differential chromatin states. Differential chromatin states are shown on the x axes (top: combinatorial state in monocytes, bottom: combinatorial state in CD4). **(b)** Expression from monocytes and CD4 cells of genes which overlap the given differential chromatin state. Numbers give the base-10 logarithm of the multiple testing corrected p-value for the expression difference using a Wilcoxon rank-sum test. The more negative the number, the more significant the difference. Genes with differential chromatin signatures (e.g. [me1/2/3+ac] in monocytes but not in CD4 T-cells) show highly significant expression differences, whereas similar signatures ([me1/2/3+ac] in both cell types) are not significantly different.

**Figure 8.**
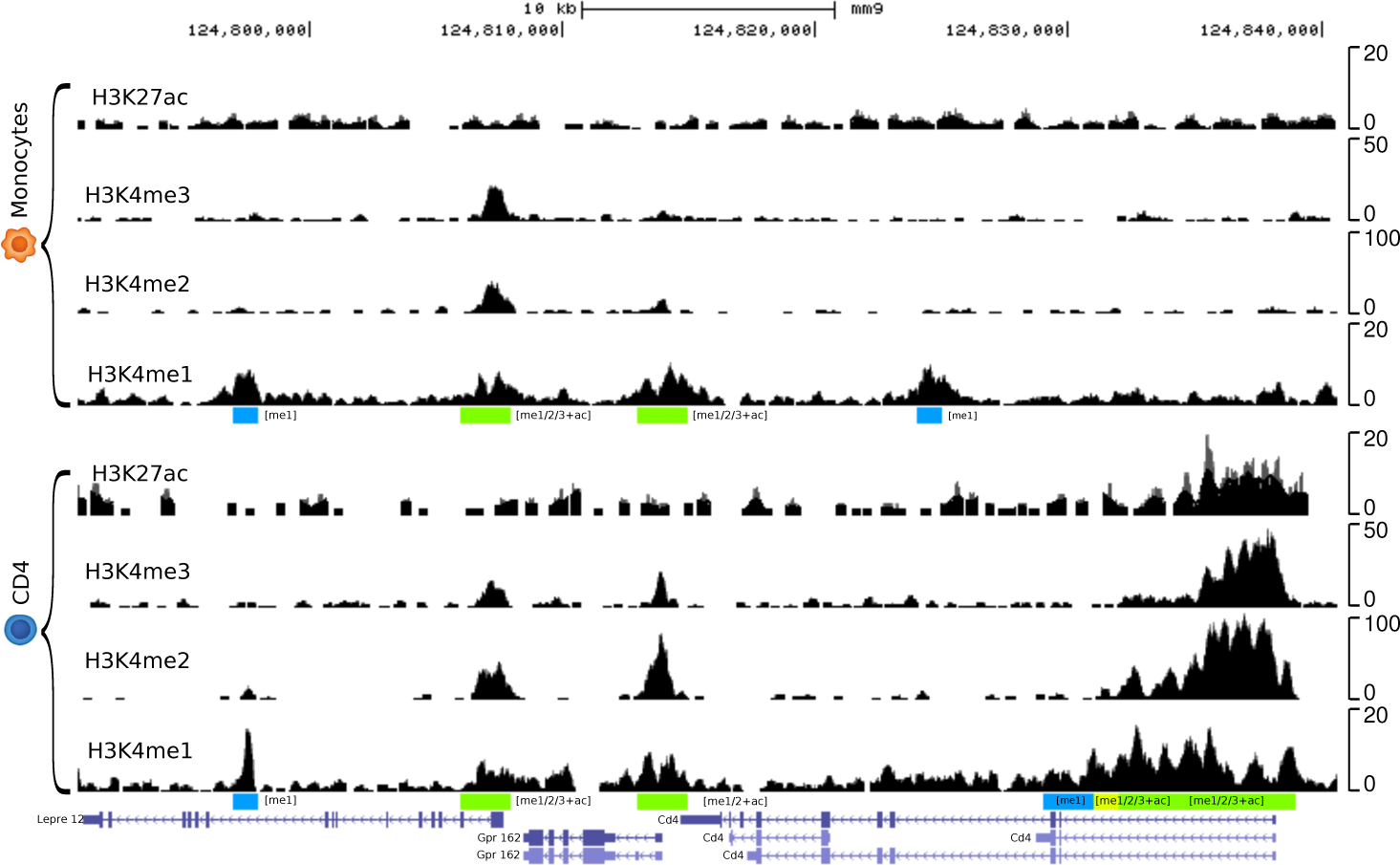
Differential chromatin signature at the Cd4 locus. Example of a differential promoter and enhancer signature at the Cd4 gene. The differential promoter signature [me1/2/3+ac] is only present in CD4 T-cells (shaded green), while the differential enhancer [me1] is present only in monocytes (shaded blue).

Again, we compared our results on the same dataset with MACS2 [33] and ChromHMM [16]. Neither method was specifically designed to deal with differences between combinatorial states, but both tools represent approaches that could have been chosen for that task in the absence of other suitable methods. For both methods, the percentage of the epigenome that was differentially modified was found to be 2.5 times higher than predicted by chromstaR, 13.02% for MACS2 and 13.59% for ChromHMM. MACS2 found most differences (3.90%) in state [me1], followed by the combination [me2+ac] (2.11%). None of these states yielded any significant enrichment in GO terms or showed correlation with expression data (Fig. S9c and Table S2). The third most frequent differential state was [me1+ac] (1.88%) and this state yielded GO term enrichments which reflect cellular identity. ChromHMM predicted two “enhancer-like states” E8 and E9 (Fig. S9b) as most differential between cell types (2.71% and 2.54%) which also showed cell type specific terms in the GO analysis (Table S3). However, expression analysis showed that ChromHMM’s most frequent differential state (CD4:E12 and Mono:E14) corresponded to proximal genes that were transcriptionally nearly inactive (Fig. S9b), which raises the question if these differential chromatin states produce cell-specific functional differences.

### Application 3: Tracking combinatorial chromatin state dynamics in time

Arguably the most challenging experimental setup is when several histone modifications have been collected for a large number of conditions, such as different cell types along a differentiation tree or different terminally differentiated tissues (Fig. S1). We consider *M* conditions with *N* histone modifications measured in each of them. This leads to *2^N^* possible combinatorial states per condition, or alternatively to *2^M^* differential states per mark across all samples. Therefore, the number of possible dynamic combinatorial chromatin states is 2*^M × N^*. For *M* × *N* ≤ *C* the whole dynamic/combinatorial chromatin landscape is treatable computationally, while for *M* × *N* > *C* the problem becomes intractable with current computational resources. The value of *C* is dependent on computational resources, genome length and bin size (see section **Limitations**).

We considered again the mouse hematopoietic data from [37], with four histone modifications (H3K4me1, H3K4me2, H3K4me3 and H3K27ac) measured in 16 different cell types during hematopoietic differentiation (stem cells, progenitor and terminally differentiated cells). We explored the chromatin dynamics during the differentiation process for every hematopoietic branch (Fig. S1a): first, long term hematopoietic stem cells (LT-HSC) are transformed into short term hematopoietic stem cells (ST-HSC) and further into multipotent progenitors (MPP). The MPP cells differentiate into the several common lineage oligopotent progenitors, giving rise to the three different hematopoietic branches (myeloid, leukocyte and erythrocyte). Finally, after another one or two stages, cells become fully differentiated at the bottom of the tree. Every branch from root to leaf consists therefore of four histone marks in five or six time points, with 2*^M × N^* = 1048576 or 16777216 possible dynamic combinatorial chromatin states, respectively. Because this number is computationally intractable, we implemented the following two-step approach for each branch: (1) for each of the four histone marks separately, we performed a multivariate differential analysis along the five or six cells in the branch, therefore assigning every bin in the genome to one of the 32 or 64 possible differential states; (2) We reconstructed the full combinatorial chromatin state dynamics by combining the differential calls of all four marks in step 1, bin by bin (Fig. S10a).

Using this two-step approach, we studied the dynamics of the inferred chromatin states over developmental time. We observed an initial increase in the frequency of the [me1] state from the LT-HSC to intermediate progenitor stages, followed by a decrease to the fully differentiated stages (Fig. S11). This decrease in [me1] was especially pronounced in the lymphoid and erythroid lineage. In the [me1/2/3+ac] signature we found a small but continuous decrease from LT-HSC to terminally differentiated stages. These observations are consistent with the view that chromatin transitions from an open configuration in multipotent cells to a closed configuration in differentiated cells. Figure S12 shows two examples of pluripotency genes, Gata2 and Cebpa, that lose their open chromatin configuration in differentiated CD4 T-cells.

We next explored the specific dynamic chromatin state transitions that occur in every region of the genome during the differentiation process. We found that the majority of all possible dynamic chromatin state transitions were not present in this system. For example, in the CD4 T-cell branch of the hematopoietic tree there are 5 developmental time points and at each stage 16 combinatorial states can be theoretically present. This leads to 16^5^ = 1048576 potential transitions between combinatorial states in this branch. However, we found only 1086 different chromatin transitions and the first most frequent 99 transitions (with frequency ≥ 0.01%) already involved 99.60% of the genome. To summarize these transitions further, we grouped them into 4 different classes: (1) “Empty” transitions, i.e. those regions that have no histone modification in any of the developmental stages. (2) “Constant” transitions, i.e. those regions that show the same (non-empty) combinatorial state in all stages of differentiation. (3) “Stage-specific” transitions, i.e. those regions that show a combinatorial state only in a subset of differentiation stages and are in the “empty” state otherwise. (4) All other transitions (see Fig. 9 for examples). In the CD4 T-cell branch, 85.98% of the genome has no measured chromatin signature in all 5 stages (class 1). The constant transitions (class 2) comprise 5.87% of the genome, stage-specific transitions 5.69% (class 3) and all other transitions 2.46% (class 4). Altogether, only 8.15% of the genome changes its chromatin state during differentiation and more than half of these changes are due to changes in the [me1] signature. This signature is highly cell type specific and gains and losses correspond to stage-specific terms in a GO analysis (Table S4) and to changes in gene expression (Fig. S13a). Among the constant transitions, regions with signature [me1/2/3+ac] mark constitutively expressed genes (Fig. S13a). Therefore we expect those regions to be enriched with housekeeping functions, which is confirmed by the GO analysis (Table S5).

**Figure 9.**
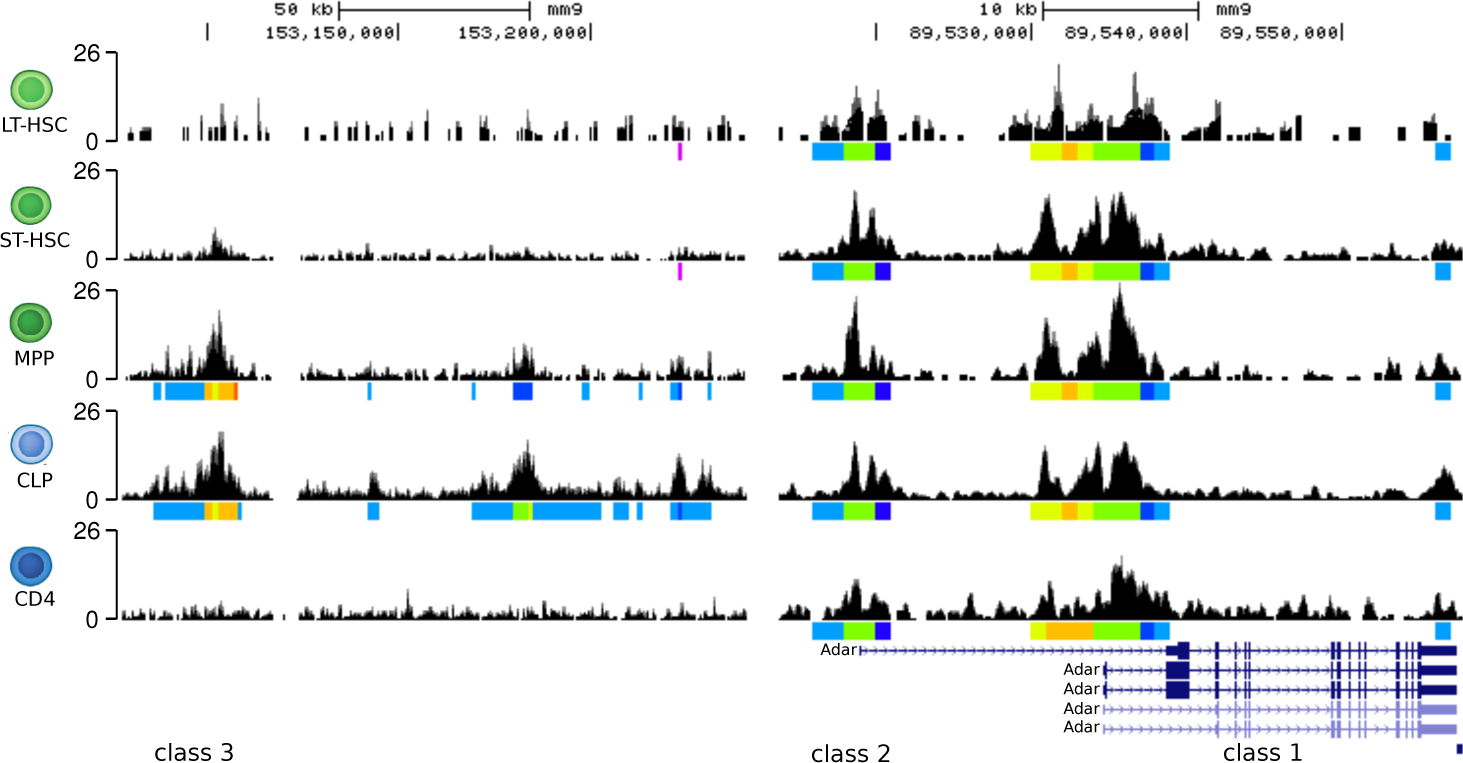
Examples of different classes of transitions (shaded in red): “Empty” (class 1), “constant” (class 2) and “stage-specific” transitions (class 3). Only the H3K4me1 tracks are shown. Combinatorial chromatin states as obtained by chromstaR are shown below the H3K4me1 tracks as colored bars

We compared our results on the CD4 T-cell branch with MACS2 [33] and ChromHMM [16]. Strikingly, MACS2 found 34470 different chromatin state transitions, with the most frequent 330 (with frequency ≥ 0.01%) covering 94.47% of the genome. This large number is expected since MACS2 is a univariate peak caller and not designed for differential analysis. Furthermore, this dataset represents a differential analysis not between two cell types, but between five different cell types and thus boundary effects (false positives, e.g. falsely detected differences) are extremely likely. This interpretation is supported by the expression data, which could not find clear expression differences for the most frequent differentially modified regions (Fig. S9c). Also the GO analysis could not identify any significant GO terms. ChromHMM found 38288 different state transitions of which the first 656 cover 91.21% of the genome. This large number of transitions is dependent on the number of states that are used to train ChromHMM, since extra states will artificially inflate the number of chromatin state transitions. However, consistent with the chromstaR predictions, ChromHMM predicts many stage-specific enhancer (state E15 and E16) and constant promoter (state E9) regions among the most frequent transitions. The expression profiles associated with those transitions show the expected behaviour (Fig. S13b).

### Limitations and Solutions

The number of possible combinatorial states for *N* ChIP-seq experiments is *2*^N^, meaning that for each additional ChIP-seq experiment the number of combinatorial states doubles. Thus it soon becomes computationally prohibitive to consider all combinatorial states. We found that with current computational resources (Intel Xeon E5 2680v3, 24 cores @ 2.5 GHz, 128GB memory) a practical limit seems to be 256 states (= 8 experiments) with a run-time of several days for a mouse genome (≈ 2.6·10^9^bp) and a bin size of 1000bp (≈ 2.6M datapoints). We investigated several possibilities to extend the usability of chromstaR beyond this limit: (1) Calculations can be performed for each chromosome separately, and chromstaR features an option to perform this calculation in parallel. (2) For the case of one cell type or tissue where the number of measured histone modifications *N* exceeds the upper limit, chromstaR provides a strategy to artificially restrict the number of combinatorial states to any number lower than 2^N^. This strategy can yield proper results if the correct states are included, since our results have shown that the majority of combinatorial states are absent in the genome. In order to identify the states which are the most present in the genome, chromstaR ranks the combinatorial states based on their presence according to the combination of univariate results from the first step of the chromstaR pipeline. This ranking is a good approximation of the true multivariate state-distribution (Fig. S14). (3) If there are multiple marks *N* in multiple tissues *M*, and 2*^N * M^* is bigger than the maximum number of states that the algorithm can handle computationally, two strategies are possible: One can either perform a differential analysis for each mark and then reconstruct combinatorial states in a classical way (Fig. S10a) or one can perform a multivariate peak-calling of combinatorial states for each tissue and then obtain the differences by a simple comparison between tissues (Fig. S10b). Both strategies give a different perspective on the data: The former accurately identifies differences between marks, while the combinatorial states might be subject to boundary effects. The latter gives an accurate picture of the combinatorial chromatin landscape, while differences between cells might be overestimated. (4) The run-time of our algorithm scales linearly with the number of data points, and thus a strategy is to decrease the resolution, e.g. halfing the run-time by doubling the bin size.

## Discussion

Understanding how various histone modifications interact to determine cis-regulatory gene expression states is a fundamental problem in chromatin biology. It is becoming increasingly clear that certain combinatorial patterns of these modifications define discrete chromatin states along the genome. These chromatin states “encode” cell-specific transcriptional programs, and constitute funtional units that are subject to dynamic changes in response to developmental and environmental cues.

Many experimental studies have recognized this and collected ChIP-seq data for a number of histone modifications on the same or different tissue(s) as well as for several developmental time points. Integrative analyses of such datasets often present formidable bioinformatic challenges. Only a few computational methods exist that can analyze multiple histone marks simultaneously in one sample and cluster them into a finite number of chromatin states [16, 17, 18, 19, 20, 21, 22]. Interestingly, these methods often demand that the user specifies the number of chromatin states beforehand. We find this problematic because this number is often a desired output of the analysis rather than an input. Indeed, the true number of distinct chromatin states in the genomes of various species is subject to debate. In *D. melanogaster* nine chromatin states have been reported [45], while in *A. thaliana* four main states were found [46]. In human, Ernst et al. found 51 states in human T-cells [47]. The Roadmap Consortium reported 15 to 18 states [10]. It remains unclear whether these differences reflect species divergence at the level of chromatin organization, or whether they are due to differences in the assessed chromatin marks and bioinformatic treatment of the data. Without a formal computational framework for defining chromatin states these two possibilities cannot be confidently distinguished.

While multivariate methods such as ChromHMM provide possible computational solutions to such questions, these methods employ probablistic chromatin state definitions that are not always readily interpretable. A probalistic interpretation means that different combinatorial histone modification patterns can be simultaneously part of different underlying chromatin states. However, it is not immediately obvious whether the underlying chromatin state are biologically distinct or if they are only statistical entities that are otherwise biologically redundant. Identifying such redundancies is not easy, because of a lack of rules to decide whether two or more chromatin states can or cannot be considered to be equivalent. Such decisions require extensive manual curation of the output, and often presuppose the kind of biological knowledge that one wishes to obtain from the data in the first place.

In contrast to this probabilistic state definition, chromstaR outputs discrete chromatin states that are defined on the basis of the presence/absence of various histone modifications. That is, with *N* histone modifications, it infers all *2^N^* combinatorial chromatin states (Fig. 1a). This interpretation makes it easy to relate the inferred chromatin states back to the underlying histone modification patterns and thus fashions a direct mechanistic link between chromatin structure and function. Moreover, chromstaR’s discrete state definition also provides an unbiased picture of the genome-wide frequency of various chromatin states and allows for easy genome-wide summary statistics. For instance, in our analysis of four histone modifications in mouse embryonic stem cells we found that only 7 of the 16 possible states covered almost 100% of the genome, and for the human Hippocampus with seven modifications only 21 of the 128 possible combinatorial states already covered 99% of the genome. Even more extreme, when analyzing 26 marks in a lung fibroblast cell line, we found that only 0.02% of all possible combinatorial states explain 95% of the genome. This striking sparsity in the combinatorial code is interesting and points at certain biochemical contraints that determine which histone modifications can or cannot co-occur at a genomic locus. Clearly, the genome-wide frequency of inferred combinatorial chromatin states depends on the number and the type of different histone modifications that are used in the analysis. Future studies should systematically investigate the dependency of the number of chromatin states on factors such as number and type of measured histone marks, resolution, organism etc.

By treating discrete combinatorial chromatin states as units of analysis chromstaR can also easily track chromatin state dynamics across cell types or developmental time points. In that respect chromstaR is unique as no other methdods exist to date that can peform a similar task. To illustrate this we have analyzed four different histone modification in 5 different cell types that are part of the mouse T-cell differentiation pathway. Of the 1048576 combinatorial state transitions, we find that only 99 comprise over 99.60% of the genome. Again, the sparsity in state transitions shows that a few key transitions define the developmental trajectory of T-cell differentiation. One notable transition is the gain or loss of state [me1] near promoters. We note that this state means that only H3K4me1 is present at a locus and no other marks. This is not the same as tracking H3K4me1 modification by itself as this latter mark can appear in a number of different, and often funtionally distinct, chromatin states such as [me1+ac], [me1/2+ac], [me1/2/3]. Hence, focusing on H3K4me1 alone would tag other chromatin state changes that may not be fully informative about T-cell differentiation.

## Conclusions

chromstaR is a computational algorithm that can identify discrete chromatin states from multiple ChIP-seq experiments and detect combinatorial state differences betweeen cell-types and/or developmental time points. By defining chromatin states in terms of the presence and absence of combinatorial histone modification patterns, it provides an intuitive way to understand genome regulation in terms of chromatin composition at a locus. chromstaR can be used for the annotation of reference epigenomes as well as for annotation of chromatin state transitions in well-described developmental systems. The algorithm is written in C++ and runs in the popular R computing environment. It therefore combines computational speed with the extensive bioinformatic toolsets available through Bioconductor [48, 49]. chromstaR is freely available at http://bioconductor.org/packages/chromstaR/ and features a collection of downstream analysis functions.

## Methods

### Model Specification

The construction of the multivariate Hidden Markov Model can be divided in two steps (Fig. 2). In the first step, we fit a univariate Hidden Markov Model to each individual ChIP-seq sample. The obtained parameters of the mixture distributions are then used in the second step to construct the multivariate emission distributions. Finally, the multivariate Hidden Markov Model is fitted to the (combined) ChIP-seq samples. The following sections describe the two steps in detail.

#### Univariate Hidden Markov Model

For each individual ChIP-seq sample, we partition the genome into *T* nonoverlapping, equally sized bins. We count the number of aligned reads (regardless of strand) that overlap any given bin *t* and denote this read count with *x_t_*. Following others [29, 30], we model the distribution of the read counts *x* with a two-component mixture of (zero-inflated) negative binomial distributions. In our case, the first component describes the *unmodified* regions and is modeled by a zero-inflated negative binomial distribution. The second component describes the *modified* regions and is modeled by a negative binomial distribution. Furthermore, for computational efficiency, we split the first component into the zero-inflation and the negative binomial distribution [31]. Our univariate Hidden Markov Model has thus three states *i: zero-inflation, unmodified* and *modified*. We write the probability of observing a given read count as

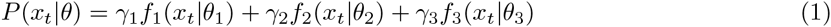
where *γ_i_* are the mixing weights and *θ_i_* are the component density parameters. The emission distribution of state 1 is defined as

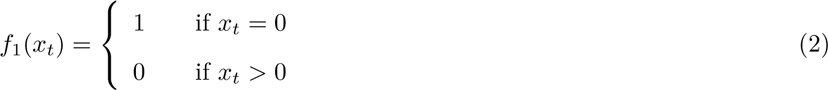
and the emission distributions of state 2 and 3 are defined as

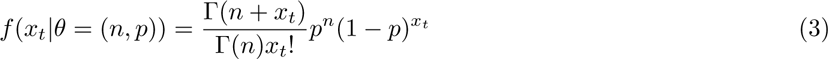
where Γ denotes the Gamma function and *p* and *n* denote the probability and dispersion parameter of the negative binomial distribution, respectively.

We use the Baum-Welch algorithm [50] to obtain the best fit for the distribution parameter estimates, transition probabilities and posterior probabilities of being in a given state. We call a bin *modified* if the posterior probability of being in that state is > 0.5 and *unmodified* otherwise.

#### Multivariate Hidden Markov Model

Given *N* individual ChIP-sep samples with states *unmodified* and *modified*, the number of possible combinatorial states is 2*^N^*. Let x*_t_* be the vector of *N* read counts for the *t*-th bin. The probability of observing a random vector x_t_ can be written as a mixture distribution of 2*^N^* components:

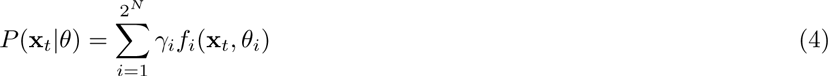

Again, the *γ_i_* denote the mixing weights and *θ_i_* denote the component density parameters for each component *i*. We assume that the marginal densities of the multivariate count distributions *f_i_* are given by the univariate distributions described in the previous section. A convenient way to construct a multivariate distribution from known marginal (univariate) distributions is copula theory [32, 51].

Under the assumption of a Gaussian copula, the multivariate emission density for combinatorial state *i* can be written as

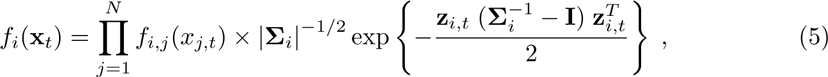

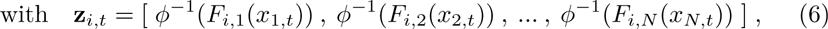
where *f_i,j_* are the marginal density functions for combinatorial state *i* and *Σ_i_* is the correlation matrix between the transformed read counts *z_i,t_* = *ϕ*^−1^(*F_i_*(*x_t_*)). The cumulative distribution function (CDF) of *f_i,j_* is denoted by *F_i,j_*, while *ϕ^−1^* denotes the inverse of the CDF of a standard normal [52].

The correlation matrix *Σ_i_* for a given multivariate (combinatorial) state *i* is computed as follows: From the combination of univariate state calls (*unmodified* or *modified*) of all samples, we pick those bins that show combinatorial state *i*. The read counts *x_t∈i_* in those bins are transformed to z*_t∈i_* using equation 6 and the correlation matrix *Σ_i_* is calculated from the transformed read counts.

Similarly to the univariate Hidden Markov Model, we use the Baum-Welch algorithm to obtain the best fit for the transition probabilities and posterior probabilities of being in a given state. However, the emission densities remain fixed in the multivariate case. We assign a combinatorial state to each bin by maximizing over the posterior probabilities. We found it useful to transform posterior probabilities for each combinatorial state (“posteriors-per-state”) into posterior probabilites of peak calls for each experiment (“posteriors-per-mark”), such that a cutoff can be applied instead of maximizing over the posteriors. We found that out algorithm had a very high sensitivity for detecting peaks when maximizing over the “posteriorsper-state”. To increase specificity, a strict cutoff (e.g. 0.99) can be applied on the “posteriors-per-mark”.

#### Integration of chromatin input experiments

Chromatin input experiments serve as controls for bias in chromatin fragmentation and variations in sequencing efficiency [53]. We optionally integrate this information by modifying the vector of read counts that serves as the observable in the Hidden Markov Model. Let x be the vector of read counts along the genome for the ChlP-seq experiment, and y be the vector of read counts for the input experiment. Let furthermore y*_P_* be the vector y without zero read counts. In a first step, we null regions with artificially high read counts, e.g. repetitive regions around centromers, by setting *x_t_* = 0 for all bins *t* where *y_t_* >= *c*_0_. *c*_0_ is defined as the 99.9% quantile of y*_P_*. In a second step, we calculate a corrected read count x′ as

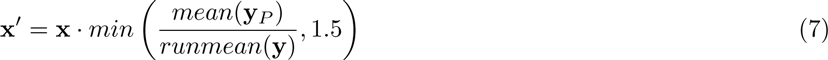
where *runmean()* calculates a running mean of 15 bins. This operation modifies the read count x in such a way that x′ is decreased in bins which have more than average counts in the input and increased in bins that have less than average counts in the input.

#### Inclusion of replicates

The chromstaR formalism offers an elegant way to include replicates. For a single ChIP-seq experiment, there are two states - unmodified (background) and modified (peaks). For an arbitrary number of *N* experiments, there are thus 2*^N^* combinatorial states. The same is true for an arbitrary number of replicates *R*, which would yield 2*^R^* combinatorial states. However, in the case of replicates, the number of states can be fixed to 2, such that all replicates are forced to have the same state (e.g. either peak or background). Treating replicates in this way allows to find the most likely state for each position considering information from all replicates without prior merging.

#### Univariate approximation of multivariate state distribution

chromstaR offers the possibility to restrict the number of combinatorial states to any number lower than 2*^N^*, where *N* is the number of ChIP-seq experiments. Because the first step of the chromstaR workflow is a univariate peak calling, we can combine those peak calls into combinatorial states and use their ranking to determine which states to use for the multivariate peak-calling. Because most systems seem to be sparse in their combinatorial patterns, i.e. do not utilize the full combinatorial state space, it is often not necessary to run the multivariate part with all 2*^N^* combinations. For instance, for the human Hippocampus tissue with seven marks, running the multivariate with only 30 instead of 128 states recovers 98.2% of correct state assignments compared to the full 128 state model, and choosing 60 instead of 128 states recovers already 99.5% of correct state assignments compared to the full 128 state model (Fig. S14).

### Data Acquisition

ChIP-seq data for the hematopoietic system (GSE60103) was downloaded from the Gene Expression Omnibus (GEO) and aligned to mouse reference mm9 following the procedure in [37] with bowtie2 (version 2.2.3) [54], keeping only reads that mapped to a unique location. The number of identical reads at each genomic position was restricted to 3. For the expression analysis, we used the provided RNA-seq data (GSE60101). We normalized the read counts by transcript length and scaled them to 1M reads. To reduce the effect of extreme expression values, we applied an arc-sinh transformation on the data.

For the Hippocampus dataset, bed-files with aligned reads were downloaded from ftp://ftp.genboree.org/EpigenomeAtlas/Current-Release/sample-experiment/Brain_Hippocampus_Middle/ for donors number 112 and 149 and seven histone marks H3K27ac, H3K27me3, H3K36me3, H3K4me1, H3K4me3, H3K9me3 and H3K9ac.

Bed-files with aligned reads for the IMR90 dataset were downloaded from ftp://ftp.genboree.org/EpigenomeAtlas/Current-Release/sample-experiment/IMR90_Cell_Line for 26 histone marks (H2AK5ac, H2BK120ac, H2BK12ac, H2BK15ac, H2BK20ac, H2BK5ac, H3K14ac, H3K18ac, H3K23ac, H3K27ac, H3K27me3, H3K36me3, H3K4ac, H3K4me1, H3K4me2, H3K4me3, H3K56ac, H3K79me1, H3K79me2, H3K9ac, H3K9me1, H3K9me3, H4K20me1, H4K5ac, H4K8ac, H4K91ac).

### Enrichment profiles around TSS

We calculated sensitivity (recall), precision and F1-score for the detection of expressed TSS based on the following assumptions: True positives are expressed TSS which are called into the promoter state ([me1/2/3+ac] for chromstaR, E7 and E9 for ChromHMM, [me1/3] and [me3] for MACS2, see Fig. 4). False negatives are expressed TSS which are not assigned into the promoter state. True negatives are non-expressed TSS which are not assigned into the promoter state. False positives are non-expressed TSS which are assigned the promoter state. We found that chromstaR has a higher sensitivity than the other methods and a lower precision. The F1-score is highest for chromstaR (Table 2).

## Availability of data and materials

ChIP-seq data for the hematopoietic system (GSE60103) was downloaded from the Gene Expression Omnibus. Bed-files for Hippocampus tissue were downloaded from ftp://ftp.genboree.org/EpigenomeAtlas/Current-Release/sample-experiment/ for donors number 112 and 149. Bed-files for IMR90 cell line were downloaded from ftp://ftp.genboree.org/EpigenomeAtlas/Current-Release/sample-experiment/IMR90_Cell_Line. *chromstaR* if available under the Artistic-2.0 license and can be downloaded from http://bioconductor.org/packages/chromstaR/.

## Author’s contributions

MCT and FJ designed the research. AT, MCT, MAN and MH developed the algorithm. AT analyzed the data. AT, MCT and FJ wrote the manuscript.

## Acknowledgements

We thank G. de Haan and JJ. Schuringa for their comments. MCT acknowledges support from the Netherlands Organization for Scientific Research and from a University of Groningen Rosalind Franklin Fellowship. FJ was supported by the Technische Universitat Munchen – Institute for Advanced Study, funded by the German Excellence Initiative and the European Union Seventh Framework Programme under grant agreement #291763. MH acknowledges support by the Federal Ministry of Education and Research (BMBF, Germany) in the project eMed:symAtrial (01ZX1408D).

